# Below-ground ants follow pheromones more quickly under dark conditions, but pheromones do not affect decision accuracy nor aggression

**DOI:** 10.64898/2026.02.16.706118

**Authors:** Patrick Krapf, Michael Mitschke, Anna Lenninger, Nico Völlenklee, Tomer J. Czaczkes, Birgit C. Schlick-Steiner, Florian M. Steiner

**Affiliations:** Molecular Ecology Group, Department of Ecology, Universität Innsbruck, Technikerstr. 25, Innsbruck 6020, Austria; Organismal and Evolutionary Biology Research Programme, University of Helsinki, Helsinki, 00560, Finland; Institute for Biodiversity and Ecosystem Dynamics, University of Amsterdam, Amsterdam, 1090 GE, Netherlands; Department of Zoology, Stockholm University, Svante Arrhenius väg 6A, 114 18 Stockholm, Sweden; Centre for Palaeogenetics, Stockholm University, Svante Arrhenius väg 20C, 114 18 Stockholm, Sweden; Animal Comparative Economics Laboratory, Chair of Zoology and Evolutionary Ecology, Institute of Biology, Universität Regensburg, Universitätsstr. 31, 93053 Regensburg, Germany; Animal Cognition and Ecology Laboratory, Applied Zoology and Animal Ecology group, Institute of Biology, Freie Universität Berlin, Albrecht-Thaer-Weg 6, 14195 Berlin, Germany

**Author notes:** shared first authors (Patrick Krapf, Michael Mitschke). shared last authors (Birgit C. Schlick-Steiner, Florian M. Steiner).

**Keywords:** *Tetramorium alpestre*, Y-maze, speed-accuracy trade off, aggression assay, artificial trail pheromones, fossorial

## Abstract

Communication allows organisms to quickly convey information vital for survival or fitness. Chemical communication and speed-accuracy trade-offs are ubiquitous in animal decision making. Most studies have used species which forage mainly above-ground species, tested in an epigean setting, but it remains unclear whether below-ground species behave similarly. Here, we use the below-ground ant *Tetramorium alpestre* to assess the efficacy of above- vs. below-ground mazes, the accuracy of decisions when using natural vs. artificial pheromones, the presence of a speed-accuracy trade-off, and the pheromones’ effect on aggression. Ants decided more quickly under below-ground than above-ground conditions, indicating they may be distracted by above-ground stimuli. Ants followed both natural and artificial trails but in direct competition preferred artificial trails, likely due to a higher pheromone concentration. Surprisingly, no speed-accuracy trade off was observed during path decision-making. Lastly, population origin but not pheromones affected if and how aggression occurred in presence of trail and home-range marking pheromones. We argue that the design of behavioural tests should match the lifestyle of the focal organism. We further speculate that speed-accuracy trade-offs likely are highly species- and context-specific and other factors besides chemicals seem important to trigger aggression, at least in this species.

## 1. Introduction

Communication is key, regardless of whether it occurs within or between species. It allows animals to convey valuable information such as territory borders, the location of food, and the presence of predators and prey (1). Conveying such information quickly and reliably can be vital for animals’ fitness and survival (2). Insects, and in particular, eusocial insects such as ants have evolved a sophisticated recognition and communication system using chemicals (3–5).

Ants use different chemicals for different purposes. For example, cuticular hydrocarbons (CHCs) are long-chained alkanes, alkenes, and methyl-branched alkanes (6) present on insects’ cuticle. The primary function of CHCs is protection against desiccation (6), but ants also use qualitative and quantitative differences in CHCs to recognise species and species (6). Another type of chemicals for communication are pheromones, which are small volatile proteins and peptides. Ants release such pheromones to transmit information (7, 8) via various glands (9), which can cause alarm or attraction effects in individuals of the same and different species (8).

Many ant species use trail pheromones to convey information on food sources (5, 9). In more detail, after finding a valuable food source, ants leave a pheromone trail on the way back to their colony (9). After being released, such pheromones can dissipate within a few minutes but also last up to hours (7, 10). Ants thus need to constantly re-apply pheromones to let their nestmates know that food is still available. Such trail pheromones can also be created artificially, for example, by synthesizing the known trail pheromone (9) or by extracting pheromones from the gaster of ants in a solvent (11, 12). Artificial trail pheromones allow us to address questions regarding information use strategies, such as when animals choose to copy others rather than innovate or following their own memories (13), or what information content is important for assessing the value of social information (14). For such assays, T- and Y-mazes are frequently used (15), in which the test pheromone is applied on one side of the maze and a control or different pheromone on the other side. Many such studies focused on above-ground foraging species such as *Cataglyphis niger* (16), *Euprenolepis procera* (11), *Lasius niger* (17), and *Linepithema humile* (18). However, our understanding of whether ant species living nearly entirely below ground perform similarly in such tests is limited. To our knowledge, only one fossorial ant – *Lasius flavus* – has been studied in such a situation, but in this experiment the ants were forced to forage in an above-ground (epigeal) setting (13).

Here, we used the below-ground-living ant *Tetramorium alpestre* (19, 20), which is a behaviourally plastic ant displaying peaceful to aggressive behaviour (21, 22). Specifically, we 1) test the efficacy of a closed dark Y-maze simulating below-ground conditions by comparing it with one simulating above-ground conditions, 2) compare natural pheromone trails with artificial ones, 3) assess the accuracy of these decisions, and 4) test whether trail as well as home-range marking pheromones affect aggression. We investigated decision accuracy, the time until decisions were reached, a potential speed-accuracy trade off (i.e., an ant decides quickly but then follows the side without pheromones vs. take longer to gather information and decides to follow the side with pheromones), as well as the influence of pheromones on aggression levels (Tab. 1).

**Tab. 1.**
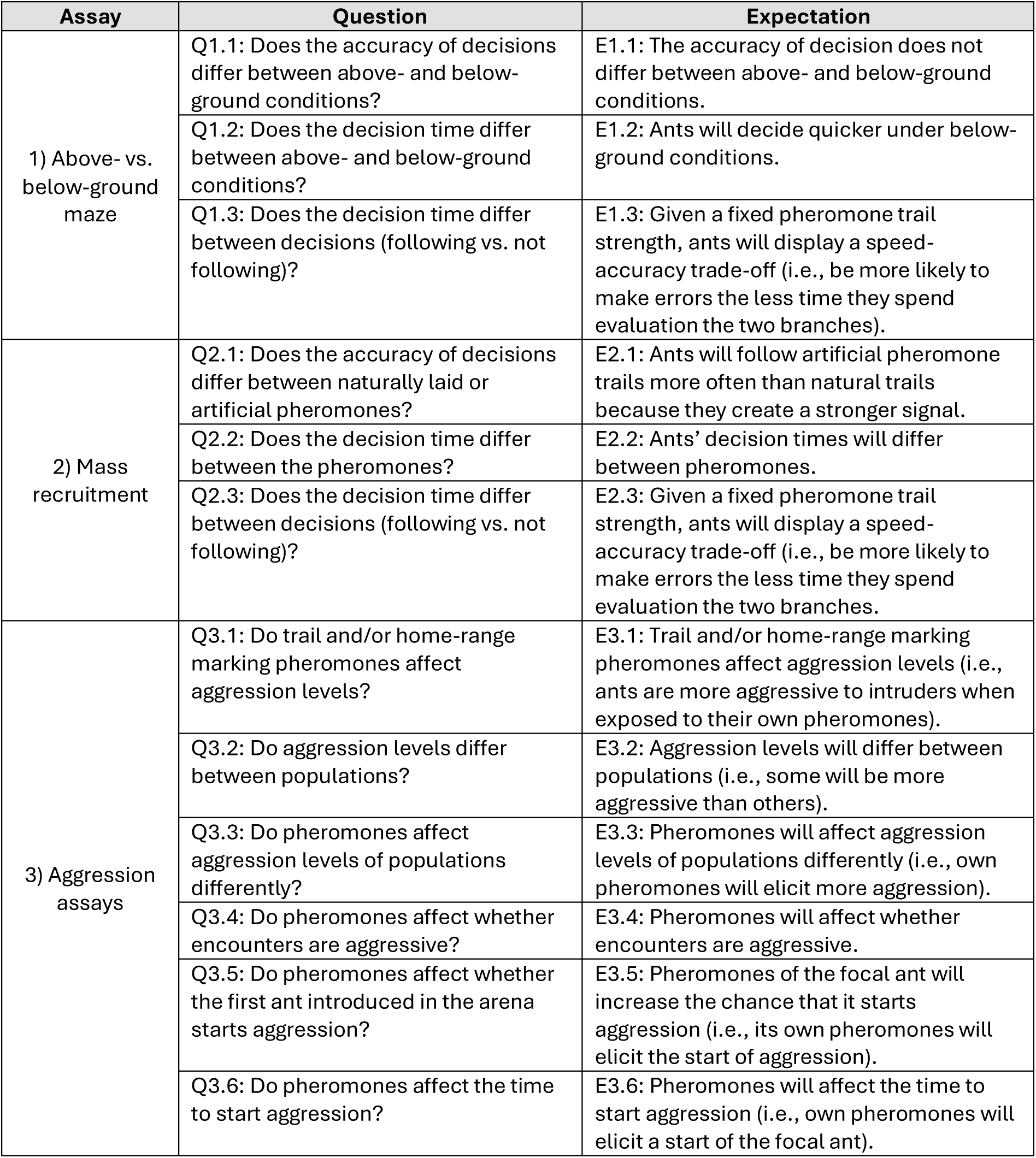
Questions and expectations for each assay.

We found that ants decided more quickly in the maze simulating below-ground (i.e., ‘dark’) conditions than in above-ground conditions. However, the decisions or time needed until ants reached a decision did not differ in their accuracy between the mazes. Further, ants followed both natural and artificially created trail pheromones more often than a control, and ants followed artificial trails more often and quickly than naturally created ones. In none of the tests, a speed-accuracy trade off was observed, but ants needed slightly longer to decide to follow a pheromone than to follow a control. Lastly, pheromones did not affect the level of aggression, whether ants displayed aggression at all, whether ants started aggression, or the time until aggression started. In contrast, aggression levels differed across populations. Aggression in this ant thus seems to be influenced by other factors than pheromones.

## 2. Material and methods

### 2.1. Colony collection and maintenance

In September 2021, 13 colonies of the ant species *Tetramorium alpestre* were collected from four populations. Seven colonies were collected from two populations in South Tyrol, Italy (Population 1 at Jaufenpass with four colonies and Population 4 at Penser Joch with three colonies; Fig. 1A; Tab. S1) and six colonies from two populations in Tyrol, Austria (Population 2 at Kuehtai and Population 3 at Hahntennjoch with three colonies each). One additional colony was collected at Jaufenpass because the number of individuals in one colony was too low for the experiment. Based on previous studies, both populations in Italy were assumed to be aggressive (21, 23), while both populations in Austria were assumed to be peaceful (21, 24).

**Fig. 1.**
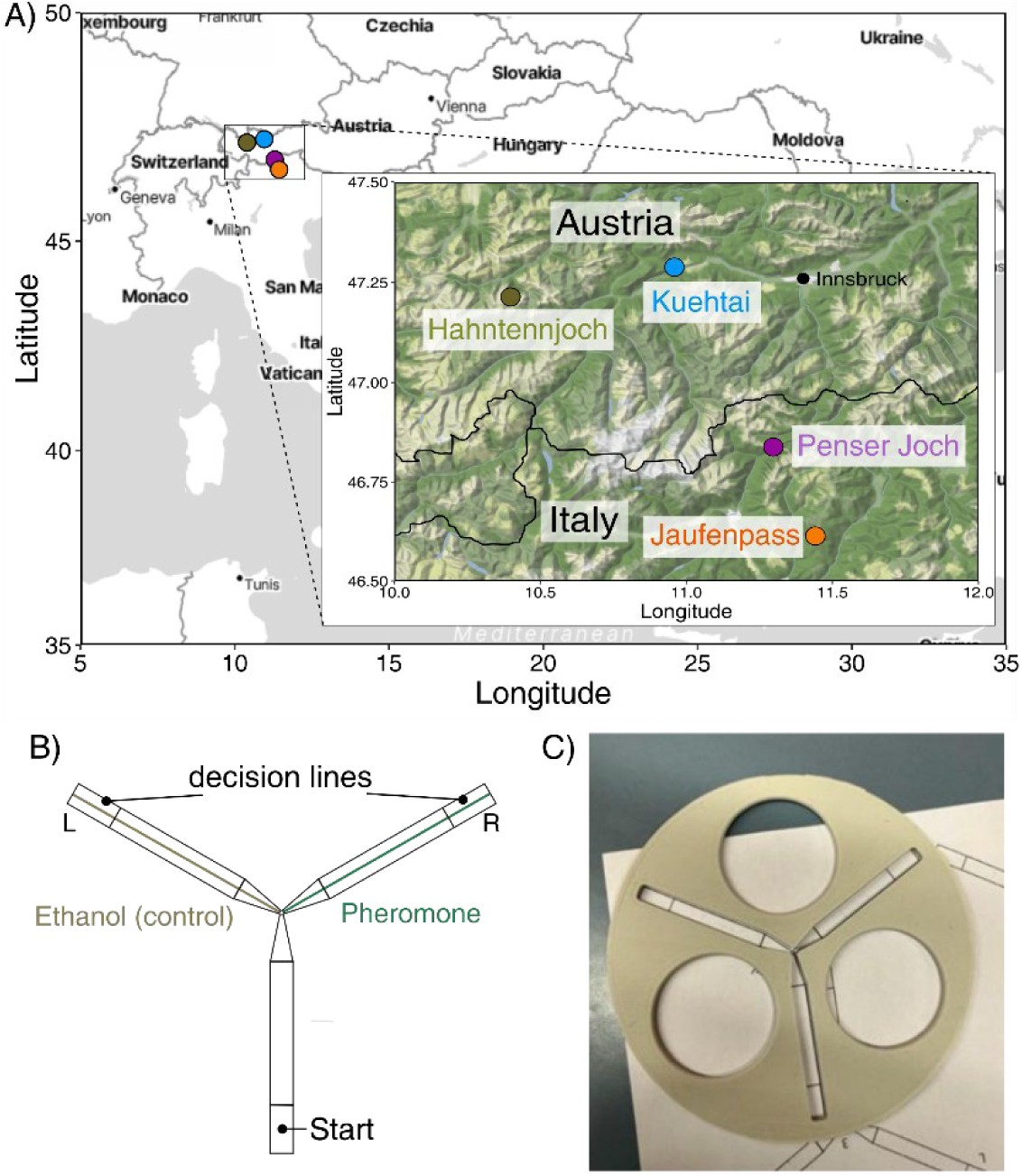
Map and schematic overview of the maze. **A)** Sampling map with the four populations used. **B)** Schematic overview of the Y-maze used and C) a photograph of the Y-maze without the red overlay used to simulate below-ground conditions.

Approximately 300 workers were collected from each colony using a polypropylene aspirator. The workers remained in the aspirator in a cooling box until they were transferred to polypropylene boxes ("nest box", 10.5 × 10.5 cm) on the same day at a laboratory at the University of Innsbruck. Small holes were drilled in the lid of the nest boxes to ensure ventilation. The walls of the nest boxes were coated with Fluon (GP1, De Monchy International BV, Rotterdam, Netherlands) to prevent workers from escaping. The floor of each nest box was made with plaster, and each box contained a small, darkened collection tube (Qiagen collection tube, 2 ml) filled with water and a ball of cotton wool as a source of water. Diluted sugar water in a collection tube and protein and sugar powder (AntStore, Reith im Alpbachtal, Austria) were available as food *ad libitum*. The nest boxes were kept in a climate cabinet (MIR-254,158 Panasonic, Etten Leur, Netherlands) at the following day-night rhythm and respective temperatures: 16 h light at 22 °C and 8 h darkness at 15 °C, as well as with approx. 60-70% relative air humidity, simulating their below-ground lifestyle (19). This day-night and temperature regime simulated a prolonged summer which allowed us to conduct the assays until November 2021. Colonies were acclimatised to laboratory conditions for two weeks before conducting experiments. For the pheromone assays, workers were starved for five days prior to the experiment, to set a high and standardised foraging motivation (25).

### 2.2. Pheromone extracts

#### Artificial trails: Extracts from the gaster

A gaster extract (henceforth “*GAS*”) was prepared for each colony. Five workers each were sacrificed in Eppendorf tubes filled with 96% ethanol p.a. briefly before the pheromone extraction. The gaster of each ant was cut off with a sterilised razor blade, and all five gasters were transferred to a conical 5 ml Eppendorf tube filled with 2.5 ml with 96% ethanol p.a. The gasters were pestled with an autoclaved pestle for approx. five minutes to extract pheromones from all glands.

#### Natural trails formed via mass recruitment

To create natural trails, colonies were first starved for five days. Each colony was then connected to an above-ground Y-maze setup with paper strips (Video 1). Workers were allowed to forage on sugar water. When returning from the food source, workers laid pheromones on the paper overlays (henceforth “*NAT*”). The number of workers following the first laid pheromone trail increased gradually (i.e., mass recruitment). The mass recruitment was terminated as soon as the number of ants feeding on the sugar water started decreasing. The papers were then retrieved, immediately cut into smaller pieces, and frozen at -20 °C until the start of the mass recruitment trails, detailed below.

### 2.3. Pheromone assays

#### 2.3.1. Blind observations

Before the start of all pheromone and aggression assays, interim colony IDs were assigned by a person not involved in the project. This means all observers (N_AboveVsBelow_ = 2; N_MassRecruitment_ = 2; N_AggressionAssay_ = 2) were blind to the colony’s origin to minimise any potential observer bias. However, the observers knew the interim colony ID during handling.

#### 2.3.2. Above- vs. below-ground maze

Testing whether workers differed in their decision (following or not following) and the time to reach a decision was achieved using two different set-ups: one maze simulating above-ground conditions and one simulating below-ground conditions. The above-ground maze consisted of a 3D-printed (ecoPLA; 3DJAKE, Paldau, Austria) customised Y-maze with a 120° bifurcation, following (15). The dark maze simulating below-ground foraging behaviour was a newly designed inverted version of the above-ground model, representing a ‘tunnel’ (Fig. 1B-C). After transferring a worker into the maze, it was covered with a transparent lid covered with red transparent foil. Ants only negligibly perceive the red-colour spectrum, which thus simulates below-ground conditions (26), allowing us to observe them. Through this procedure, ants were also forced to remain in the device during the experiment. Both mazes used standard printer paper as substrate, which was replaced after each experiment to remove natural pheromone laid by the workers. Each paper arm additionally contained printed black lines, which in the assembled maze represented one “start line” and two “decision lines” (Fig. 1B).

Before transferring the workers to the Y-mazes, either 2.5 μl of a trail pheromone or a control (96% ethanol p.a., henceforth “*CON*”) was applied as a thin line onto one of the two arms using a 10 µl-syringe (Hamilton, Bonaduz, Switzerland). After applying the trail pheromones, the syringe was cleaned with 96% ethanol p.a. at least twice. The side onto which the trail pheromone or ethanol was applied was alternated between experiments. This procedure minimised any effects of potential side bias, which are known to affect ants (27). After applying the trail pheromone or control, the ethanol was left to evaporate for at least one minute before starting the experiment.

The experiment started by transferring an individual worker onto the Y-maze using soft forceps. For the below-ground assay, the experiment began closing the maze with the red cover. Whenever a worker crossed the starting line or one of the decision lines, we recorded which line it was and the time until the decision line was crossed. For each worker, we also recorded the colony origin, the side on which the pheromones and the control were applied, and the time the worker needed to cross the start line. After the worker had crossed the decision line (regardless of the side), the assay was finished. The experiment was terminated after three minutes, when the worker had not crossed the start line and after five minutes, when it had crossed the start line but not the decision line. The experiments were carried out until 20 final decisions had been recorded for all colonies. After each individual experiment, the maze, forceps, and lids were thoroughly cleaned with ethanol. After a completed experiment, the ants were transferred into a new nest box to prevent re-using of workers. Workers from four colonies (1I, 2C, 3F, 4K) were tested on two consecutive days in October 2021. During the assays, each of the two observers tested all colonies by assaying two colonies, with each on one side above-ground and below-ground and then switched colonies and pheromones.

#### 2.3.3. Mass recruitment

This assay tested whether ants followed i) the natural trail pheromone *NAT* or ii) the artificial *GAS* extract more often and more quickly than ethanol as control *CON*, and if workers followed iii) *GAS* more often and more quickly than *NAT*. The assay was conducted on two consecutive days using four colonies, two of them, colonies 1B and 2E on the first day, and two, colonies 3L and A4 on the consecutive day. Each of the two observers assayed one colony for 15 experiments, namely five experiments using paper overlays with *NAT* vs. *CON*, five experiments using paper overlays with *NAT* vs. *GAS*, and five experiments using paper overlays with *GAS* vs. *CON*.

#### 2.3.4. Statistical analyses of the above-ground and below-ground maze and mass recruitment assays

For the statistical analyses, the final decision of the ant was used and termed “following” or “not following” depending on whether the ant followed the pheromone trail or not, respectively. If a worker did not reach a decision, this was noted as “no decision”. All statistical analyses were conducted in R v.4.3.0 (28) using RStudio v2024.12.1 (29). Before analysing the data, we checked for each assay whether a) the left and right arm of the Y-maze had been used equally often using a binomial test, b) workers did not follow one side more often than expected by chance using a binomial test, and c) there is an observer bias (e.g., only noting the right side as result) using a Chi-square test with Yates’ continuity correction. In the mass recruitment assay, natural pheromone trails (*NAT*) were used, which may weaken over time. For these, we tested whether a possible pheromone evaporation over time resulted in a change in the decision by conducting a Chi-square test with Yates’ continuity correction on the decision of ants between the pheromone trails. We further checked whether decision times were normally distributed for the assays mentioned above using a Shapiro-Wilk test in R.

To answer questions Q1.1, Q2.1, and Q3.1, whether ants followed pheromone trails under above- vs. below-ground conditions or a pheromone type more often than another setup or pheromone, a Chi-square test with Yates’ continuity correction was used with all data as well as binomial tests for each setup or pheromone separately. To answer questions Q1.2, Q2.2, and Q3.2, whether ants followed a setup or pheromone more quickly than another one, we tested whether data were normally distributed using a Shapiro-Wilk test. Most data were not normally distributed, so a Mann-Whitney-U test was used to compare the arithmetic means of decision times between the setups or pheromones. To test whether the proportions of ants differed that either followed *GAS* or *NAT*, a 2-sample test for equality of proportions with continuity correction was conducted (‘prop.test’ in R).

### 2.4. One-on-one aggression assays

#### 2.4.1. Preparation

To test if pheromones influence aggressive behaviour, one-on-one encounters were conducted in a flat-bottomed glass arena with paper discs (1.4 cm diameter; standard printer paper) at the bottom of the arena. In experiment (a), 1 μl of 96% ethanol p.a. was pipetted onto paper discs as a control (*CON*) to assess if alcohol affects aggression tests. In experiments (b) and (c), untreated paper discs were placed into the nest box of each focal colony for at least 24 hours (*24H*). The paper discs were thereby coated with the colony odour of the focal colony, such as territorial and home-range marking pheromones (30). In these two experiments, we further pipetted 1 μl of 96% ethanol p.a. onto the paper discs to make it comparable with the control and experiments (d) and (e). In experiments (d) and (e), 1 μl of the gaster extract *GAS* was pipetted of the respective colony onto the paper disc.

Thirty minutes before the start of the aggression tests, the colonies were transferred from the climate cabinets to the laboratory to bring them to room temperature. Each aggression test was filmed using a hand camera (Handycam HDR-XR 155; HDR-PJ810E, Sony, Tokyo, Japan; or camcorder HC-V777, Panasonic, Osaka, Japan).

#### 2.4.2. Experimental setup

Aggression tests were conducted within each population using three colonies, yielding three combinations per population. For each combination, we replicated the above-mentioned experiments (a-e) five times (Replicates 1-5) to capture possible behavioural variation (31). In total, 25 tests were carried out per colony combination. Workers had access to an *ad libitum* amount of food and water before being used in the aggression assays.

After the respective pheromone had been applied to the paper disc, the ethanol was allowed to evaporate for at least four minutes before using the paper discs. The paper discs were then placed on the bottom of the glass arena. Then, two naïve workers (one from the focal colony and one from a different colony of the same population) were randomly selected and placed in the glass arena with the paper disc. The first worker was transferred using tweezers and the video-recording was started. A few seconds later, the second worker was added. This allowed us to distinguish between the two workers in the recordings. The inner walls of the flat-bottomed test tubes were coated with Fluon (GP1, De Monchy International BV, Rotterdam, Netherlands) to prevent the ants from escaping.

To minimise the bias of circadian rhythm in the aggression assays, the first replicates of the experiments (a-e) were conducted in a consecutive order, starting with experiment (a). Then, the second replicates of experiments (a-e) starting with experiment (b), etc. were conducted. To further minimise the bias of workers that were introduced first into the glass arena in experiment (a), we placed an ant from one colony into the arena in replicate 1 and the other ant first in replicate 2, thus alternating which ant was introduced first. In the other experiments (b-e), the ant belonging to the colony paper disc *24H* or *GAS* extract was always added first. This procedure simulated that the respective ant was in its colony and should recognise the introduced ant as an intruder or friend. This also led to a doubled number of assays with the pheromones *GAS* and *24H* than with the control *CON*. Ants from the within-population tests were sacrificed by freezing. Additionally, we conducted within-colony aggression assays to test if the workers’ behaviour was peaceful, which we expected (not shown).

#### 2.4.3. Statistical analysis

The behaviour of each worker was recorded individually for each second and assigned a scoring scale ranging from -4 to 5: -4 = trophallaxis, -3 = mutual grooming, -2 = antennating, -1 = being next to each other without touching, 0 = ignoring, 1 = avoiding, 2 = mandible threatening, 3 = fighting without flexing the gaster, 4 = fighting with flexing the gaster, 5 = killing (32). Based on the behavioural data, a behavioural index *AI* (33) was calculated for each worker and combination using the formula

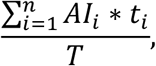

where *AI*_*i*_ and *t*_*i*_ are the aggression index and the time of each behaviour and *T* is the total duration of the behavioural video. The mean over the five replicates was calculated separately for each experiment (a-e).

To assess the impact of the pheromones and population origin on the behaviour, addressing Q3.1 and Q3.2, a pairwise Mann-Whitney-U test was conducted for the behavioural means for each pheromone and population separately (Bonferroni-Holm corrected for multiple testing). Addressing Q3.3, the impact of the pheromones was assessed separately within each population using a pairwise Mann-Whitney-U test with Bonferroni-Holm correction.

For the remaining three questions (Q3.4-Q3.6), whether pheromones elicit aggression and affect the time to start aggression, we selected only encounters in which ants were aggressive (behaviour index higher than zero indicating aggression (32)). Because of different numbers of videos for each pheromone, relative proportions of aggressive workers or workers starting aggression were calculated for each pheromone. To test if the focal pheromone elicits aggression (Q3.4), we first tested whether more encounters (i.e., relative proportions) were aggressive in one of the three pheromones using a three-sample test for equality of proportions with continuity correction. To test whether the pheromones affect whether an ant starts aggression (Q3.5), a three-sample test for equality of proportions with continuity correction was conducted. For these workers, the focal pheromone had been applied on the paper discs. Our rationale here is that this simulates the nearby presence of nestmates. Lastly, we tested whether the pheromones elicit workers to start aggression more quickly in one of the three groups (Q3.6) by comparing the time when ants started aggression across the three pheromones used by conducting a pairwise Mann-Whitney-U test with Bonferroni-Holm correction.

## 3. Results

### 3.1. Descriptive statistics

In total, 404 pheromone assays were performed in the mazes. The right and left side of the Y-Maze were used similarly often in all assays, so no bias by applying the pheromone more often to a specific side of the maze (Tab. S2) was introduced. Further, we did not observe a side preference in the workers nor an observer bias of the person recording the data (Tab. S2).

### 3.2. Comparing above- vs. below-ground mazes

To compare the above- vs. below-ground maze, four colonies (1I, 2C, 3F, and 4K) were used and *GAS* extracts or ethanol as control *CON* were tested in 184 tests. Of these, 24 were finished without any decision and were not included in the analysis. In the analysis, 160 pheromone assays were used. The decision times were not normally distributed under either condition (Tab. S3).

In both above-ground and below-ground conditions, ants followed the arms with *GAS* extracts significantly equally often (Q1.1; Tab. S3; Fig. 2A; Pearson’s Chi-Square Test for count data; Comparing *following* between above- and below-ground, χ² = 0.070, df = 1, p-value = 0.792) and significantly more often than the control arm with ethanol (Binomial tests; above-ground: *following* = 73; *not following* = 7; Binomial test, probability = 0.91, p-value <0.001; below-ground: *following* = 71; *not following* = 9; Binomial test, probability = 0.89, p-value <0.001). Ants decided significantly more quickly under below-ground than above-ground conditions (Q1.2), being on average 68.89 seconds faster (Mann-Whitney-U test, Mean decision times above-ground: 122.05 s, below-ground: 53.16 s, Mann-Whitney-U, W = 5322, p-value <0.001, Tab. S3, Fig. 2B). In the above-ground maze, ants often walked back and forth or explored different parts of the maze (e.g., climbing on the underside of the maze). No speed-accuracy trade-off was observed neither in the above- or the below-ground maze (Q1.3). Under above-ground conditions, ants took longer, but not significantly so, in deciding to follow a *GAS*-scented arm than to follow a control arm (Mean decision time *following*: 125.33 s; *not following*: 87.86 s, W = 191.00, p-value = 0.276; Tab. S3; Fig. 2C). Under dark conditions, ants followed a *GAS*-scented arm almost as quickly as following a control arm (Mean decision time *following*: 52.28 s, *not following,* 60.11 s; W = 406.5, p-value = 0.188; Tab. S3).

**Fig. 2.**
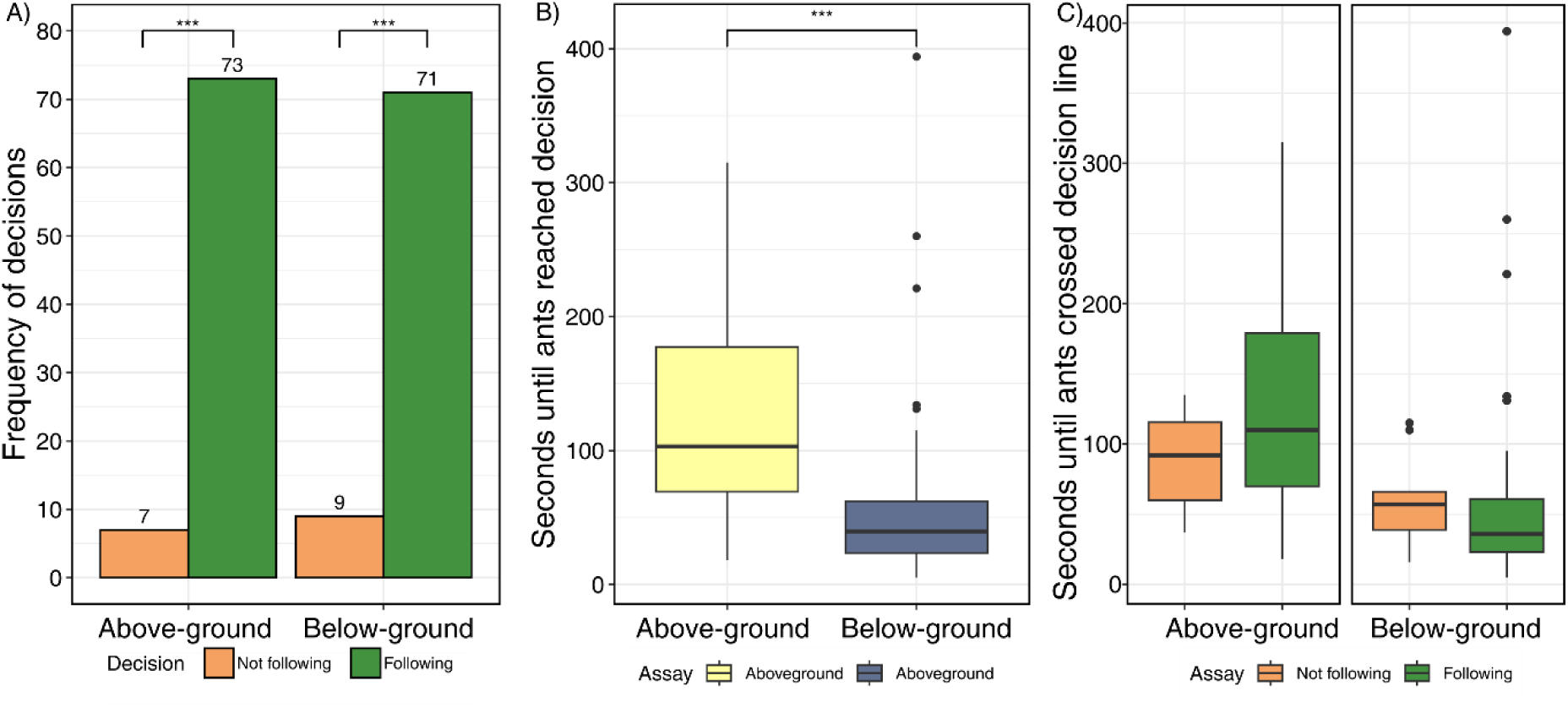
Decision frequency, decision time, and speed-accuracy trade off displayed for the above- and below-ground assays. **A)** Decision frequency of ants *Following* and *Not following* the pheromone trail. Note that the frequencies do not reach 100% as we excluded ants that did not reach a decision. **B)** Time until ants reached a decision under both setups. **C)** Decision times in seconds displayed for ants *Following* and *Not following* the pheromone trail under both setups. Asterisks indicate statistical significance with ***: p-value <0.001.

### 3.3. Mass recruitment

In the mass recruitment assays, 269 tests in the maze were conducted using natural pheromone trails *NAT* as well as *GAS* extracts (for details, see Materials and Methods and Supp Video 1). Of the 269 tests, 24 did not yield a decision and were excluded from the subsequent analyses. For the remaining 245 tests, the paper overlays used earlier or later in the assays did not affect how ants decided (Pearson’s Chi-square test, χ²<0.001, df = 1, p-value = 0.996). The time until ants reached a decision was not normally distributed (Tab. S4), but the time did not differ significantly between the papers used (Mean decision time: *paper used earlier*: 72.22 s; *paper used later*: 78.81 s; Mann-Whitney U test, W = 2799.5, p-value = 0.558).

Ants followed both natural and artificial trails more often than ethanol as control, but in the direct comparison, ants followed artificial trails more often than natural trails (Q2.1). Overall, there was a difference between the frequency of ants following the pheromones (Pearson’s Chi-square test, χ²= 55.96, df = 2, p-value <0.001), which we further investigated by conducting three comparisons. In the i) *NAT vs. CON assay*, ants followed the natural pheromone trail significantly more often than the control arm (*NAT* = 50; *CON* = 30; Binomial test, probability = 0.63, p-value = 0.033; Fig. 3A).In the ii) *GAS vs. CON assay*, ants followed the arm with *GAS* extracts significantly more often than the control arm (*GAS* = 64; *CON* = 16; Binomial test, probability = 0.80, p-value <0.001). In the iii) *NAT* vs. *GAS assay*, ants followed the arm with *GAS* extract significantly more often than the arm with natural pheromones (*GAS* = 65; *NAT* = 20; Binomial test, probability = 0.77, p-value <0.001). The proportions of ants following *GAS* (in *GAS* vs. *CON,* 80.0%) was also significantly higher than the proportions of ants following *NAT* (in *NAT* vs. *CON*, 62.5%; 2-sample test for equality of proportions, χ²=5.16, df = 1, p-value = 0.023).

**Fig. 3.**
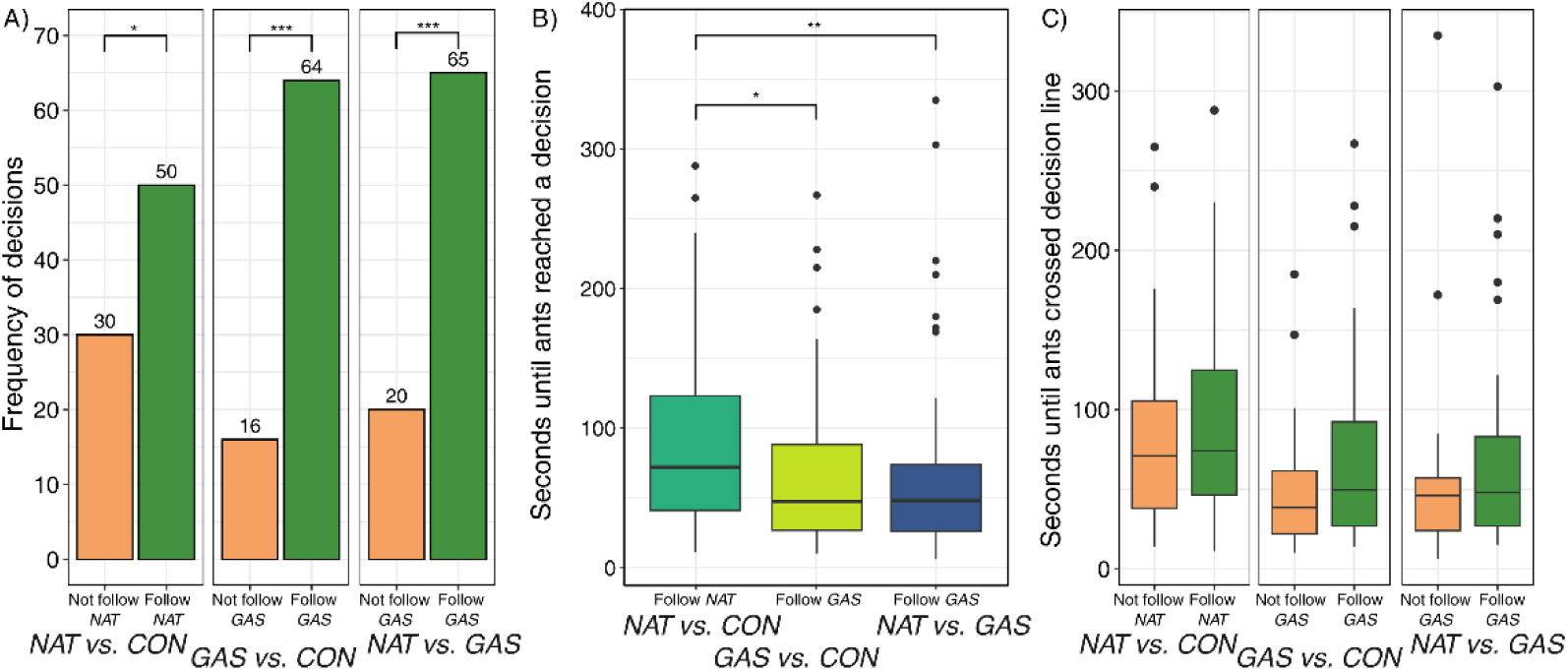
Decision frequency, decision time, and speed-accuracy trade off in the mass recruitment assays. **A)** Decision frequency of ants *Following* and *Not following* trails for i) *artificial gaster extracts GAS vs. control CON assay*, ii) *natural trail pheromones NAT vs. control CON assay*, and iii) *natural trail pheromones NAT vs. artificial gaster extracts GAS assay.* Note that the frequencies do not reach 100% as we excluded ants that did not reach a decision. **B)** Time in seconds until ants reached a decision in the three assays. **C)** Decision times in seconds displayed for ants *Following* and *Not following* the pheromone trail or control in the three assays. Asterisks indicate statistical significance with *: p-value <0.05; ***: p-value <0.001

When comparing the mean decision times between *GAS vs. CON* and *NAT vs. CON* (Q2.2), ants decided significantly more quickly when the *artificial GAS extract* was present compared with when *NAT* was present (Mean decision times *GAS vs. CON* = 65.46 s; *NAT vs. CON* = 85.31 s; Mann-Whitney-U, W = 3948.5, p-value = 0.011; Tab. S4, Fig. 3B). When comparing the *NAT vs. CON* with the *artificial GAS vs. NAT*, ants decided significantly more quickly in *artificial GAS extract vs. NAT* (Mean decision times *GAS vs. NAT* = 64.01 s; *NAT vs. CON* = 85.31 s; W = 4310, p-value = 0.003; Fig. 3B). Both values remained significant after Bonferroni-Holm correction for multiple testing. When comparing the *artificial GAS vs. CON* with *artificial GAS vs. NAT*, ants decided equally quickly (Mean decision times *GAS vs. CON* = 65.46 s; *GAS vs. NAT* = 64.01 s, W = 3345.5, p-value = 0.860).

No speed-accuracy trade-off was observed (Q2.3). The decision times of *following* vs. *not following* a pheromone trail did not differ (Fig. 3C; *NAT vs. CON*; mean decision time *following*: 79.51 s; mean decision time *not following*: 70.53 s; Mann-Whitney U test, W = 3628.5, p-value = 0.318; *GAS vs. CON*, mean decision time *following*: 67.98 s; *not following*: 55.38 s, W =597.5, p-value = 0.306; *GAS vs. NAT*, mean decision time *following*: 64.85 s; *not following*: 61.30 s, W =739.5, p-value = 0.356).

### 3.4. Aggression assays

In total, we conducted and videotaped 300 one-on-one aggression assays between different colonies (N_Workers_=600). Of the recordings, two videos were corrupted and could not be analysed, yielding a total of 298 videos from which the behaviour and interactions were evaluated according to Table 2. In addressing the first three questions (Q3.1-3), the behavioural data from all 298 aggression assays were used. In addressing the other three questions (Q3.4-6), only aggressive workers were selected that started aggression and aggressive workers starting aggression that were also introduced first resulting in 156 videos and 88 workers analysed, respectively.

**Table 2.**
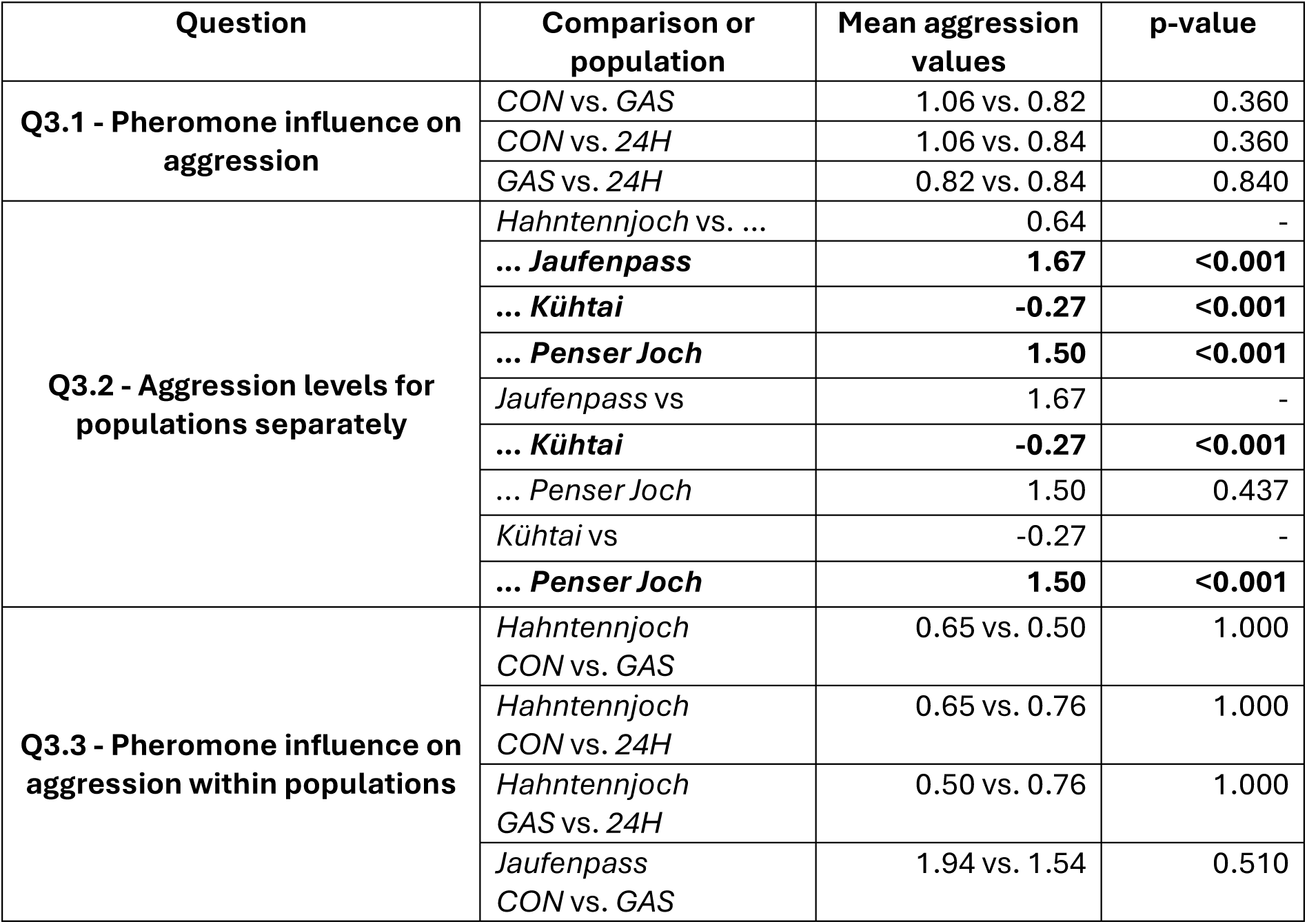
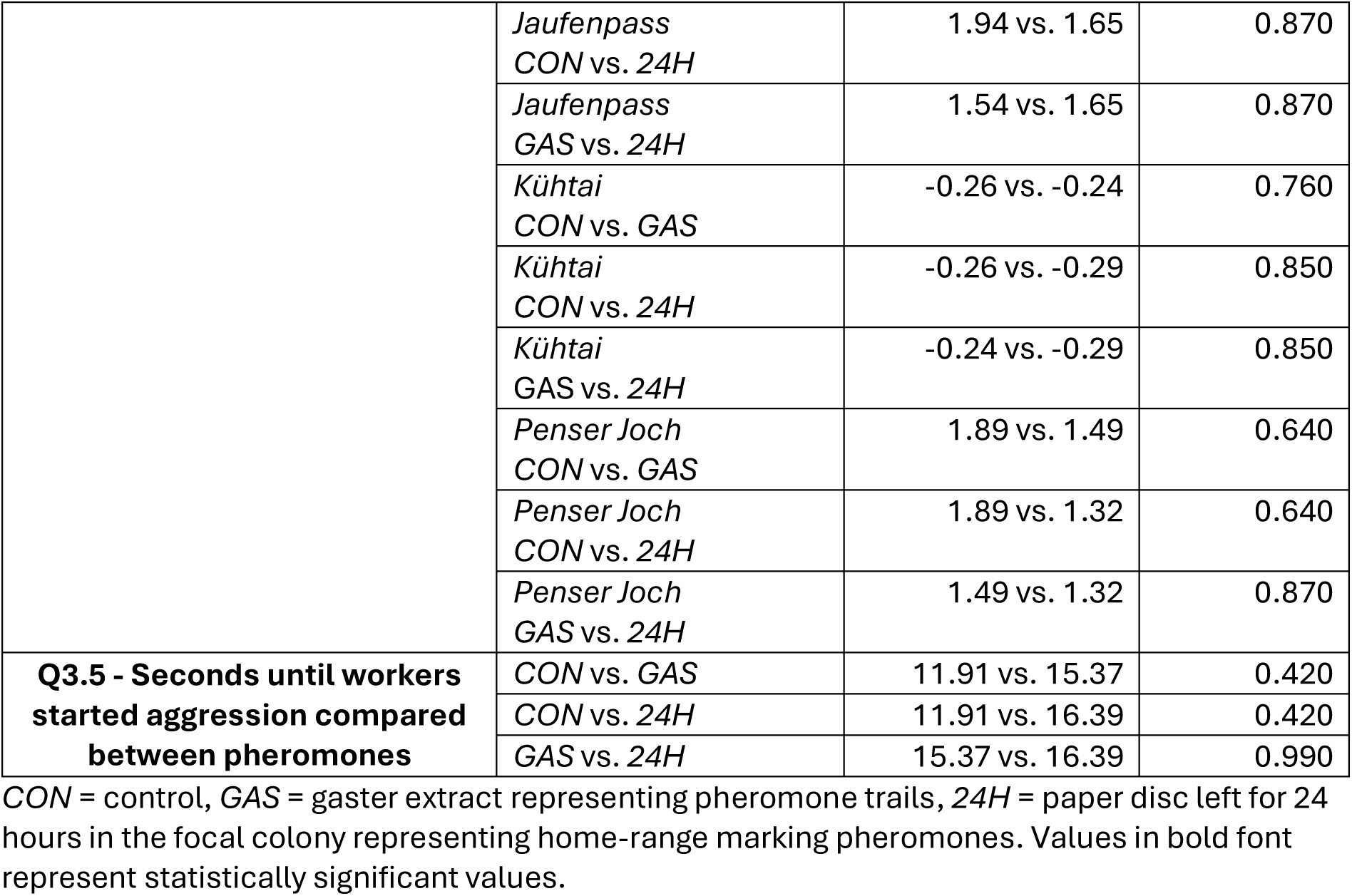
Aggression levels within and between pheromones and/or populations.

The different pheromones tested did not affect the aggression level in this ant (Q3.1). In addressing Q3.1, pheromone data across populations were combined to test whether the pheromones – *artificial GAS extract* representing trail pheromones or *24H* colony odour representing home-range marking pheromones – affected aggression compared with the control *CON*. Aggression data were not normally distributed (Tab. S5). The two pheromones and control did not impact the aggression levels when using all colonies and populations in the analysis (Fig. 4A, Tab. 2).

**Fig. 4.**
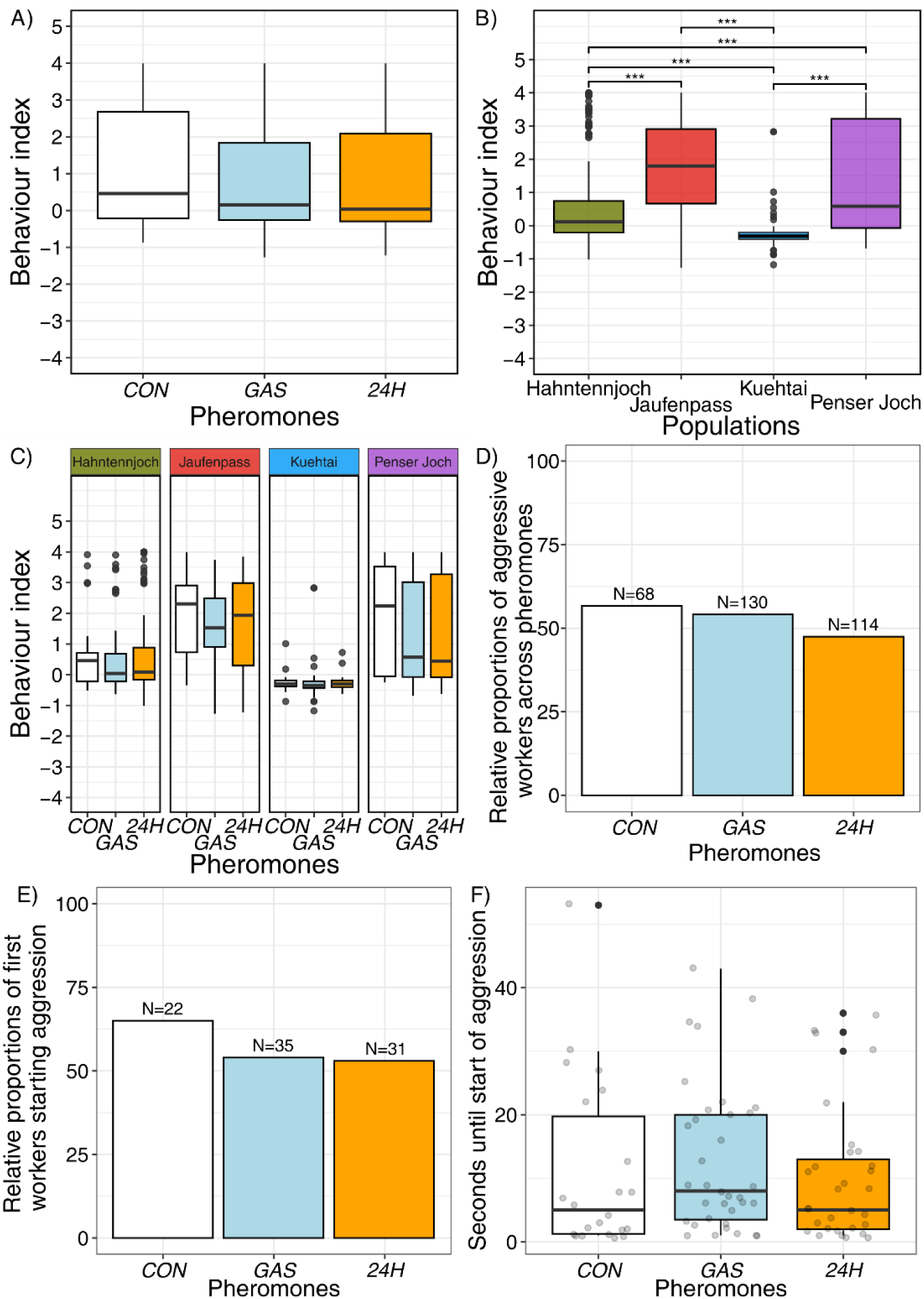
Behavioural index of the aggression assays displayed for the pheromones tested. **A)** The behavioural index is averaged over all populations and displayed separately for the three pheromones used on the paper discs. Ethanol was used as *control, CON*, *GAS* is the *artificial gaster extract* representing trail pheromones, and *24H* represents paper discs left for at least 24 hours in the focal nest box representing home-range marking pheromones. **B)** Behavioural index displayed for each population separately. **C)** Behavioural index displayed for each pheromone setup and each population separately. **D)** The relative proportions of aggressive workers are displayed for each pheromone tested. **E)** The relative percentage of workers starting aggression are displayed for each pheromone tested. We selected workers that were introduced first and also had the paper disc of the focal colony in the arena simulating a colony surrounding. In **D)** and **E)**, the numbers above the barplots represent the absolute numbers of workers tested. **F)** The seconds until workers started aggression are displayed for each pheromone tested separately. Asterisks indicate statistical significance with ***: p-value<0.001.

Aggression levels differed across but not within populations (Q3.2). Aggression data were not normally distributed (Tab. S5). Aggression levels differed significantly among all populations except Jaufenpass and Penser Joch, which were both similarly aggressive (Fig. 4B, Tab. 2; pairwise Mann-Whitney U test with Bonferroni-Holm correction, all pairwise p-values<0.001; most data were not normally distributed; Tab. S5). In detail, two populations were mainly peaceful throughout the assays (Hahntennjoch, Kuehtai), while one was always aggressive (Jaufenpass), and one (Penser Joch) displayed both peaceful and aggressive behaviours. However, pheromones did not affect aggression levels within populations (Q3.3). Different pheromones led to slight but not significant changes in the aggression levels within populations (Fig. 4C, Tab. 2).

Both pheromones did not affect whether workers were aggressive or whether workers started aggression (Q3.4, Q3.5), when testing only aggressive encounters. Regardless of the pheromone, workers were equally frequently aggressive (relative proportions of aggressive workers; *CON* = 57%, 68/120 workers, *GAS* = 54%, 130/240 workers, and *24H* colony odour 48%, 114/240 workers; χ² = 3.446, df = 2, p-value = 0.179; Fig. 4D). Also, regardless of the pheromone used, workers started aggression equally frequently when being introduced first in the arena, although the control revealed a slightly higher percentage (*CON* = 65% (22/34 workers), *GAS* = 55% (35/64), and *24H* colony odour 54% (31/57; X^2^ = 1.118, df = 2, p-value = 0.572; Fig. 4E). Similarly, pheromones did not affect the time to start aggression (Q3.6). We tested whether workers started aggression more quickly with one of the pheromones or the control. The time when aggression started was not normally distributed for each pheromone (Tab. S5). If workers were aggressive, they started aggression equally fast regardless of the pheromones or control, within 20 seconds after the onset of the aggression assays (Fig. 4F, Tab. 2).

## 4. Discussion

Social insects such as ants mainly communicate chemically (3). They use pheromones to convey information such as the trail to a food source (9). Recent studies made use of such pheromones and tested various questions using Y-mazes (15). However, most of these studies were performed with above-ground ant species. Here, we use the below-ground-living ant species *Tetramorium alpestre* to 1) compare the efficacy of a dark Y-maze simulating below-ground conditions with an above-ground maze, 2) compare natural pheromone trails with artificial ones, 3) assess the decision accuracy, and 4) test whether pheromones affect aggression.

### 4.1. Efficacy of a dark Y-maze simulating below-ground conditions

Ants of below-ground species *T. alpestre* decided more quickly when placed in mazes simulating below-ground conditions which excluded visible light by using a red lid covering the tunnel-shaped y-maze. This result is in line with our expectation because below-ground ant species are used to navigate in dark surroundings and the below-ground maze simulates such conditions. One explanation for this result could be that the pheromones are more readily encountered in a tunnel setup, because there is less evaporation and the pheromone might be easier to perceive. Another non-exclusive explanation could be that ants are less distracted under below-ground (‘dark’) than above-ground conditions because under below-ground conditions they encounter a reduced number of stimuli (e.g., optical stimuli). For instance, when using the above-ground maze with exposed arms on which the ants were placed, *T. alpestre* ants extensively explored various parts of the maze such as the underside of the arms. Further, ants may have been distracted by additional stimuli, including movements of the observers, objects on the tables, or lights in the laboratory.

Such distractions may be a general phenomenon in above-ground and thus exposed mazes. For example, in a slightly different above-ground setup, 15% of ants of the Argentine ant *L. humile* did not touch food or fell from a bridge during the assay (34). We thus find it plausible that under such conditions ants are more distracted, leading to slower decision times than in below-ground conditions. This may affect all ants, independent of their lifestyle but could be more pronounced in below-ground ants, which may encounter fewer stimuli under natural conditions.

The below-ground maze simulates a tunnel in the soil and thus represents a more natural setup. One potential drawback of such tunnels is that ants can simply walk along one side of the wall and are thus channelled into one side of the Y-maze (15, 35). However, we argue that this was unlikely to be the case here: (1) The bifurcation point in the inverted Y-maze was narrower than the tunnel, making it difficult to just follow the wall without encountering the pheromones. (2) Ants followed the arm with the pheromone more often than the arm with ethanol as control, implying that they were not simply following the wall. (3) There was no significant difference between the number of ants following a pheromone compared with the above-ground setup, confirming that the setup had no impact on the qualitative outcome of the experiment. Generally, we suggest that behavioural assays should be adjusted to the ecology and lifestyle of the study organism tested.

### 4.2. Comparing natural pheromone trails with artificial ones

We demonstrated, for the first time, that *T. alpestre* conducts mass recruitment (Supp. Video 1). Mass recruitment – when one ant finds a food source, deposits trail pheromones on the way back to the colony and thus triggers multiple other ants to follow the trail – is a quick way to share the location of a food source. Mass recruitment has been observed in other species of the *Tetramorium caespitum* group (36) such as *T. immigrans* (37).

In our assay, mass recruitment yielded significantly strong and reliable trail pheromones as ants followed natural (*NAT*) trails more frequently than ethanol as control. At the same time, ants followed the *artificial gaster extract GAS* more frequently and more quickly than the naturally laid pheromones *NAT*. This result could be due to a higher concentration of trail pheromones present in the *artificial GAS extract* than in natural trails pheromones (9). Furthermore, natural pheromones ants can dissipate within a few minutes to hours after being released (7, 10). We controlled for this by changing the paper overlays and also found no significant differences in the accuracy of ants following the paper overlays. Nevertheless, slightly decreased trail pheromone concentrations may still have contributed to a higher proportion of ants following *GAS* than following *NAT* as well as shorter decision times in following *GAS* than following *NAT*.

Besides these factors, additional compounds dissolved in *GAS* may have resulted in more ants following the *GAS* trail. In myrmicine ants, the venom gland is usually the source of the pheromone (9). Here, we used the complete content of the gaster for pheromone extracts including the Dufour and venom glands but also the crop and haemolymph. It is thus plausible that other compounds present in the gaster additionally elicited pheromone following when using the *artificial GAS extract*.

One possibility to create artificial extracts that have a concentration as found in nature are dilution experiments (11). In such experiments, the artificial extracts are gradually diluted and compared with naturally laid trail pheromones in Y-mazes. A natural concentration is reached as soon as ants decide for either of the artificial or natural trail equally often. In future assays, such diluted extract should be used.

### 4.3. No speed-accuracy trade-off in this ant

*Tetramorium alpestre* ants did not display a significant speed-accuracy trade-off (decide for ‘wrong’ decisions more quickly), but they needed a slightly longer time to decide to follow a pheromone trail than to follow ethanol as control. The relatively low error rates in the frequency of correct decisions and the increased time to reach decisions suggest that *T. alpestre* ants display a slight but not significant speed-accuracy trade-off. This minor speed-accuracy trade-off could be associated with its below-ground lifestyle, where workers likely face less predators and can take more time to reach a decision without imminent consequences. In contrast, other ant species display a speed-accuracy trade-off, for example, when *Themnothorax albipennis* ants are forced to migrate to a new colony site above-ground (38) or when *Dorymyrmex tener* ants cooperatively transport things (39). Such a trade-off may be more pronounced if a higher number of ants are involved in the decision-making process. In contrast, no speed-accuracy trade off was found in nestmate recognition in the ant *Camponotus aethiops* (40). These studies as well as our study tested speed-accuracy trade off in different species and with different settings. It thus seems probable that differences in species and the study design can generate different results.

### 4.4. Pheromones did not affect aggression

Population origin but not the tested pheromones were linked to the aggression level. None of the pheromones tested (trail pheromones, home-range marking pheromones) or the control elicited a change in the aggression levels, neither among populations nor within populations. Also, roughly the same percentages of encounters were aggressive, and workers started aggression equally frequently at a similar time after the start of the assay, regardless of the pheromones used. In contrast, aggression levels differed significantly between, but not within, the populations.

These results contrast our expectations that the focal ant should be more aggressive towards an intruder and start aggression earlier. Our expectation was based on the idea that the pheromones represent the colony’s home-range marking pheromones (*24H* papers; (30)) or trail pheromone (*GAS*) and thus simulate the presence of nearby nestmates. We assumed that ants would lay home-range marking as well as trail pheromones onto the paper discs that were present in the nest boxes for at least 24 hours. However, we did not specifically observe whether ants laid pheromones onto these paper discs. This could mean that these paper discs were similar to the control and had no effect on aggression levels. Nevertheless, since we did not observe any change in the aggression levels neither with these *24H* paper discs or with the artificial gaster extracts (*GAS*), we speculate that their effects on the aggression of ants were similarly weak and negligible.

Another explanation could be that *T. alpestre* ants use gaster pheromones mainly to signal the location of food sources. Alternatively, colony pheromones may be context-dependent (41) and do not increase aggression in the artificial setting tested here. Another possibility is that other pheromones not tested here may affect the behaviour and aggression specifically more directly. For example, mandibular gland pheromones can elicit ‘alarm’ or ‘aggression’ behaviours (42, 43).

The results suggest that aggression in the ant *T. alpestre* is mainly influenced by the population origin and population-specific environmental characteristics rather than pheromones. In previous research, we found that aggression was not observed in all populations tested (21, 22), that aggression was positively linked with increased nitrogen values and temperature (21), and that aggressive populations were bolder and more explorative and risk-prone than peaceful populations (24). However, aggression was not linked to cuticular hydrocarbons nor within-colony relatedness (21). More recently, we also observed that the start of aggression is likely linked to the gut microbiome, mutations in genes, and differentially expressed genes (44). These studies demonstrated that *T. alpestre* is a largely peaceful ant with a minor proportion of populations being aggressive. They further indicated that various other factors, besides known factors such as CHCs or relatedness, can contribute to aggression in the ant *T. alpestre* (21). Here, we demonstrated that pheromones do not affect aggression in this ant. We thus speculate that peacefulness may be linked more generally to colony and population origin and population-specific environmental characteristics in mainly peaceful ants. Reduced aggression may be adaptive in this ant, and possibly others, resulting in adaptive advantages (45). For instance, several ant species are mainly peaceful (e.g. (45–48)). They are thus not injured and not lost as a workforce. In turn, they have more time and energy available for colony growth and reproduction, yielding a fitness advantage for the colony (45).

## 5. Conclusions and outlook

We found that the below-ground ant *T. alpestre* reached a decision more quickly in dark mazes simulating below-ground conditions, likely due to a reduced number of distracting stimuli. We argue that decision making, and behavioural experiments in general, should be adapted to the lifestyle of the focal organisms to yield robust results, for instance using a below-ground maze for a below-ground ant species. Interestingly, we did not observe a significant speed-accuracy trade off, although ants needed more time to decide to follow a pheromone trail than to follow a control. We speculate that such trade-offs are highly species and context specific. Also, population origin but not pheromones tested affected the aggression levels in this ant. In previous research, we have shown that various other factors besides pheromones contribute to aggression in this ant, which could also extend to other non-aggressive ant species.

In this study, we have shown that artificial gaster extracts are perceived as trail pheromones, which has been observed in other ant species (11, 12, 15, 49). Following the methods of these other studies, we have created relatively strong artificial trail pheromones that likely have a higher pheromone concentration than natural trails. In direct comparisons, this resulted in more ants following artificial trails than natural ones. We have further laid groundwork by testing different pheromone setups in a below-ground-living ant species across three assays. Future studies should repeat at least part of the assays with an increased number of above-ground and below-ground ant species to test if our findings are generalisable. It would also be interesting to test whether this ant also follows trail pheromones of closely related con-generic ant species (9) as well as more distantly-related species, to test how closely pheromone evolution matches phylogeny. Overall, our findings highlight the complex interaction of communication and behaviour in social insects. The concentration and composition of trail pheromones as well as the experimental setup can have profound impact on the results of such behavioural experiments. The design of behavioural experiments should thus match the ecology and lifestyle of the focal organism as closely as possible to generate representative results.

## Supplementary tables

**Table S1.** Colony characteristics.

| ID | Country | Location | Latitude | Longitude | Elevation (m. a.s.l.) | Expected behaviour |
| --- | --- | --- | --- | --- | --- | --- |
| 1B | Italy | Jaufenpass | 46.8380 | 11.2978 | 1990 | aggressive |
| 1G | Italy | Jaufenpass | 46.8382 | 11.2978 | 1995 | aggressive |
| 1I | Italy | Jaufenpass | 46.8382 | 11.2980 | 2000 | aggressive |
| 1M | Italy | Jaufenpass | 46.8383 | 11.2982 | 1993 | peaceful |
| 2C | Austria | Kuehtai | 47.2140 | 10.3981 | 1896 | peaceful |
| 2E | Austria | Kuehtai | 47.2193 | 11.0750 | 1810 | peaceful |
| 2J | Austria | Kuehtai | 47.2140 | 10.5977 | 1891 | peaceful |
| 3D | Austria | Hahntennjoch | 47.2881 | 10.9593 | 1895 | peaceful |
| 3F | Austria | Hahntennjoch | 47.2883 | 10.6566 | 1875 | peaceful |
| 3L | Austria | Hahntennjoch | 47.2886 | 10.6575 | 1867 | peaceful |
| 4A | Italy | Penserjoch | 46.6139 | 11.4420 | 1974 | aggressive |
| 4H | Italy | Penserjoch | 46.8146 | 11.4425 | 1974 | aggressive |
| 4K | Italy | Penserjoch | 46.8144 | 11.4424 | 1981 | aggressive |

**Table S2.** Results of testing for a bias in the pheromone placement on the Y-maze arm, ant-side preference and observers’ scoring bias.

| Experiment | Pheromone placement |  |  |  | Ant-side preference |  |  |  | Scoring bias |  |  |
| --- | --- | --- | --- | --- | --- | --- | --- | --- | --- | --- | --- |
| | L | R | Prob. | p-value | L | R | Prob. | p-value | $\chi^2$ | df | p-value |
| **Above- vs. below-ground assay** | 80 | 80 | 0.50 | 1.000 | 84 | 76 | 0.53 | 0.580 | 2.030 | 3 | 0.567 |
| **Mass recruitment** |  |  |  |  |  |  |  |  |  |  |  |
| GAS vs. CON | 40 | 40 | 0.50 | 1.000 | 46 | 34 | 0.58 | 0.219 | 0.750 | 1 | 0.387 |
| GAS vs. NAT | 44 | 41 | 0.52 | 0.828 | 35 | 50 | 0.42 | 0.128 | 0.000 | 1 | 1.000 |
| NAT vs. CON | 41 | 39 | 0.51 | 0.911 | 47 | 33 | 0.59 | 0.146 | 2.610 | 1 | 0.106 |

We tested whether we used the left and right (L and R) arms of the Y-maze similarly often for placing the pheromones and whether the ants had a side preference using binomial tests. Further, we tested whether observers introduced a bias and were scoring a side more frequently than another side using a Pearson's Chi-squared test with Yates' continuity correction. Prob. = probability, df = degrees of freedom, GAS = gaster extract, CON = ethanol as control, NAT = natural pheromone trail on paper.

**Table S3.**
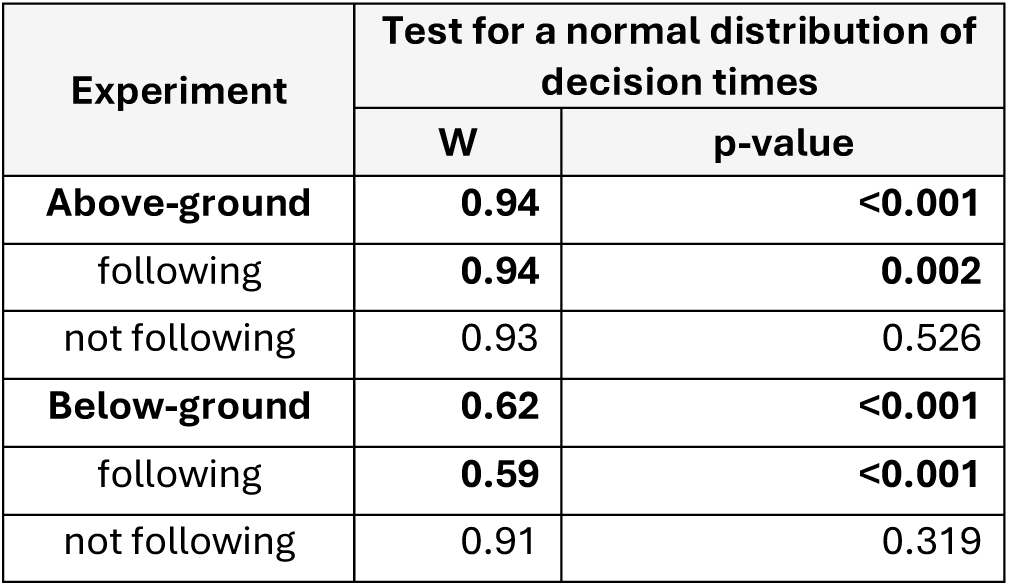
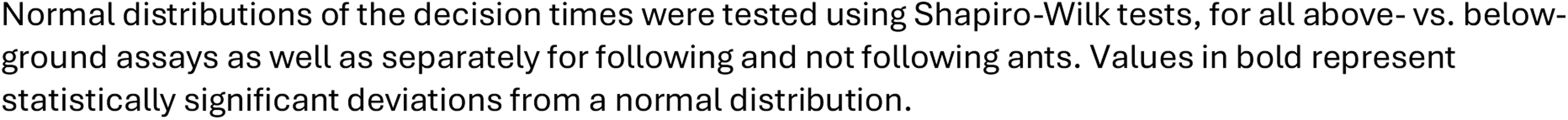
Normal-distribution test results for the decision times in the above- vs. below-ground assay.

| Experiment | Test for a normal distribution of decision times |  |
| --- | --- | --- |
|  | W | p-value |
| **Above-ground** | **0.94** | **<0.001** |
| following | **0.94** | **0.002** |
| not following | 0.93 | 0.526 |
| **Below-ground** | **0.62** | **<0.001** |
| following | **0.59** | **<0.001** |
| not following | 0.91 | 0.319 |

Normal distributions of the decision times were tested using Shapiro-Wilk tests, for all above- vs. below-ground assays as well as separately for following and not following ants. Values in bold represent statistically significant deviations from a normal distribution.

**Table S4.**
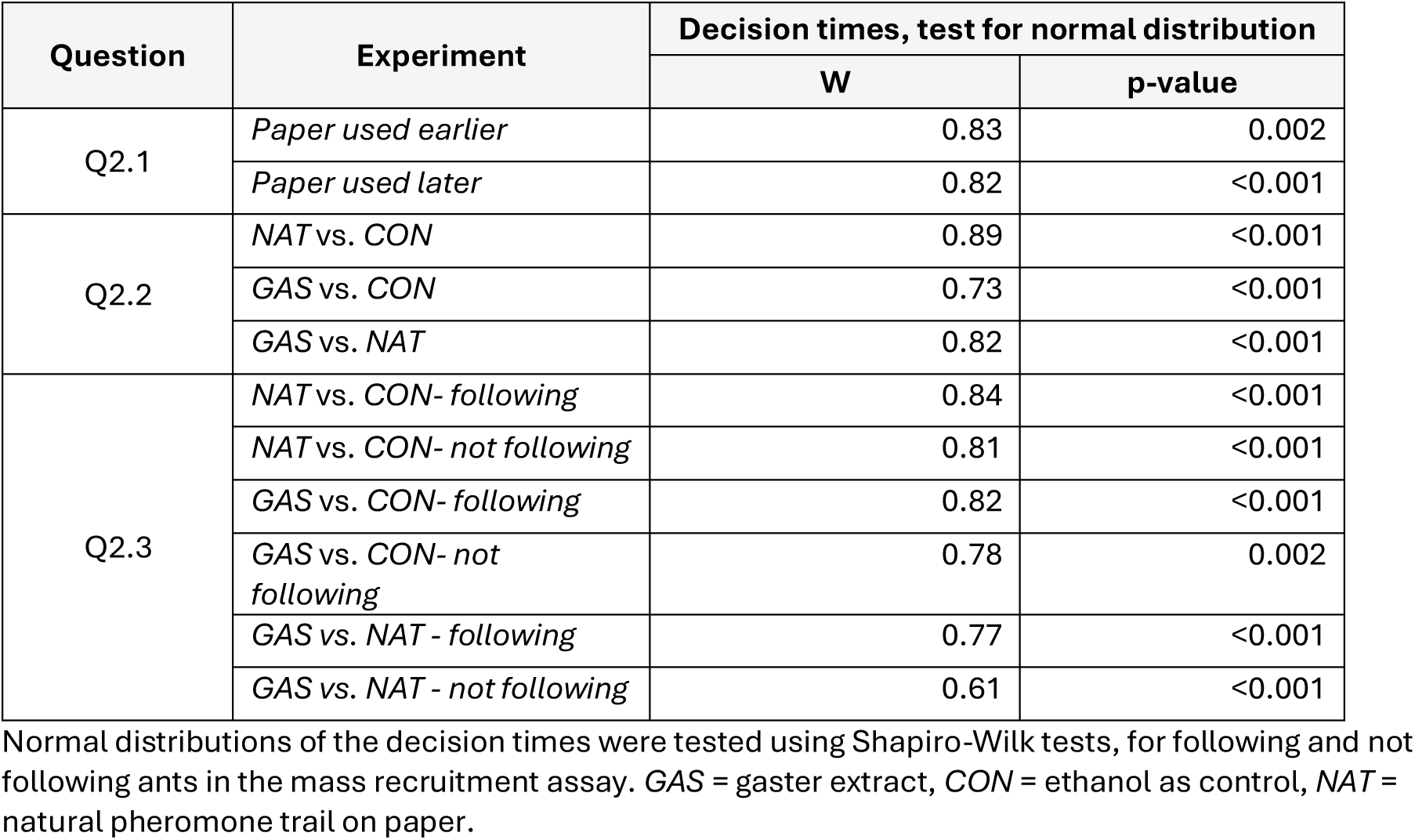
Results of the normal-distribution tests for the decision times in the mass recruitment assay.

| Question | Experiment | Decision times, test for normal distribution |  |
| --- | --- | --- | --- |
|  |  | W | p-value |
| Q2.1 | *Paper used earlier* | 0.83 | 0.002 |
|  | *Paper used later* | 0.82 | <0.001 |
| Q2.2 | NAT vs. CON | 0.89 | <0.001 |
|  | GAS vs. CON | 0.73 | <0.001 |
|  | GAS vs. NAT | 0.82 | <0.001 |
| Q2.3 | NAT vs. CON- following | 0.84 | <0.001 |
|  | NAT vs. CON- not following | 0.81 | <0.001 |
|  | GAS vs. CON- following | 0.82 | <0.001 |
|  | GAS vs. CON- not following | 0.78 | 0.002 |
|  | GAS vs. NAT - following | 0.77 | <0.001 |
|  | GAS vs. NAT - not following | 0.61 | <0.001 |

Normal distributions of the decision times were tested using Shapiro-Wilk tests, for following and not following ants in the mass recruitment assay. GAS = gaster extract, CON = ethanol as control, NAT = natural pheromone trail on paper.

**Table S5.** Normal-distribution test results for the aggression levels in the behaviour assays.

| Question | Experiment | Aggression data, test for normal distribution |  |
| --- | --- | --- | --- |
|  |  | W | p-value |
| Q3.1 | CON | 0.84 | <0.001 |
|  | GAS | 0.85 | <0.001 |
|  | 24H | 0.81 | <0.001 |
| Q3.2 | Hahntennjoch | 0.77 | <0.001 |
|  | Jaufenpass | 0.96 | <0.001 |
|  | Kühtai | 0.55 | <0.001 |
|  | Penser Joch | 0.83 | <0.001 |
| Q3.3 | Hahntennjoch CON | 0.78 | <0.001 |
|  | Hahntennjoch GAS | 0.72 | <0.001 |
|  | Hahntennjoch 24H | 0.78 | <0.001 |
|  | Jaufenpass CON | 0.93 | 0.035 |
|  | Jaufenpass GAS | 0.98 | 0.372 |
|  | Jaufenpass 24H | 0.92 | <0.001 |
|  | Kühtai CON | 0.77 | <0.001 |
|  | Kühtai GAS | 0.53 | <0.001 |
|  | Kühtai 24H | 0.82 | <0.001 |
|  | Penser Joch CON | 0.82 | <0.001 |
|  | Penser Joch GAS | 0.83 | <0.001 |
|  | Penser Joch *24H* | 0.81 | <0.001 |
| Q3.5 | *CON* | 0.64 | <0.001 |
|  | *GAS* | 0.81 | <0.001 |
|  | *24H* | 0.73 | <0.001 |

Normal distributions of the decision times were tested using Shapiro-Wilk tests, for following and not following ants in the mass recruitment assay. *CON* = ethanol as control, *GAS* = gaster extract, *24H* = paper discs present in the focal colony for at least 24 hours before the experiment.

## Ethics

This study used ants from a non-threatened ant species. No licence or permit was required to conduct assays with invertebrates including ants. Nevertheless, we followed the highest ethical standards and international law for ants collected and kept in the laboratory. We also followed two of the 3Rs Principle: i) We reduced the number of workers collected to a minimum (Reduction principle) and iii) we created the best accommodation possible for ant workers (Refinement principle). Ant colony fragments were collected in the field and maintained in the laboratory with care. Ants were provided with suitable nesting possibilities, food, and water ad libitum. After the experiments, colony fragments were kept in the laboratory and reared until their natural death.

## Data accessibility

The .STL-file to 3D-print the inverted maze as well as all data and code used in the paper will be stored in the Supplementary Materials and on GitHub upon publication.

## Declaration of AI use

AI was used to help annotate the R code. The suggested annotations were sense-checked by PK.

## Authors’ contributions

**P.K.:** Conceptualization, methodology, software, validation, formal analysis, investigation, data curation, writing – original draft, writing – review & editing, visualization; **M.M.:** Conceptualization, methodology, validation, investigation, writing – original draft, writing – review & editing; **A.L.:** Investigation, writing – original draft, writing – review & editing; **N.V.:** Investigation, writing – original draft, writing – review & editing; **T.J.C.:** Conceptualization, methodology, writing – original draft, writing – review & editing, supervision; **B.C.S.S.:** Conceptualization, methodology, resources, writing – original draft, writing – review & editing, supervision, project administration, funding acquisition; **F.M.S.:** Conceptualization, methodology, resources, writing – original draft, writing – review & editing, supervision, project administration, funding acquisition.

All authors gave final approval for publication and agreed to be held accountable for the work performed therein.

## Acknowledgements

We thank Benedikt Mader for help during colony collections, Theresia Telser for help with creating the artificial pheromones, Florian Reischer for designing the 3D sketch of the inverted Y-maze, and Philipp Bertemes, Marlene Haider, and Bob Kaufmann for help with 3D printing the Y-mazes. We thank Samuel Napetsching, Florian Reischer, Theresia Telser, and Elisabeth Zangerl for help during the pheromone assays and Evelina Krol for help with video analysis.

